# Uncompromised, multimodal, multiscale structural analysis of the hierarchically organization in mineralized tissues

**DOI:** 10.64898/2026.04.07.717027

**Authors:** Robin H.M Van der Meijden, Luco Rutten, Marit de Beer, Rona Roverts, Deniz Daviran, Judith M. Schaart, Avital Wagner, Ben Joosten, Matthijn R. Vos, Juriaan Metz, Elena Macias-Sanchez, Anat Akiva, Nico Sommerdijk

## Abstract

We present a live-to-cryo correlative imaging workflow for multiscale structural and chemical analysis of biological tissues in their near-native state. The method integrates live super-resolution fluorescence microscopy, live and cryogenic Raman spectroscopy, and targeted cryogenic focused ion beam/scanning electron microscopy, transmission electron microscopy, electron tomography, energy dispersive X-ray spectroscopy, and electron diffraction. This approach enables precise 3D targeting and nanoscale imaging of selected regions across four orders of magnitude in spatial resolution, while preserving ultrastructure and chemical composition. Using regenerating zebrafish scales as a benchmark, we visualize collagen fibril orientation, local matrix density, and mineral composition within the extracellular matrix. We identify a plywood-like architecture of unmineralized collagen with orientation-independent density variation, and reveal curved, acidic phosphate-rich mineral platelets aligned with collagen fibrils. This workflow establishes a generalizable strategy for comprehensive 3D correlative analysis of hybrid tissues, and opens new opportunities for studying native structure–function relationships at the interface of biology and materials science.

## Introduction

Understanding the structure and (bio)chemistry of hierarchically organized biomaterials is essential for deciphering how tissues develop, function, and respond to changes^1^. Such understanding requires multiscale imaging techniques that can bridge molecular composition, cellular structure, and tissue-scale architecture in three dimensions. Yet, correlating high-resolution structural and chemical information across biologically relevant length scales remains a major imaging challenge.

This challenge is especially pronounced in mineralizing tissues such as bones, teeth and seashells, as well as in calcifying soft tissues, such as arterial walls or heart valves, where hybrid structures composed of soft organic matrices and hard inorganic phases must be imaged without introducing deformation or chemical artifacts^2^. Existing imaging methods either lack the spatial resolution, compromise chemical integrity^3^, or fail to provide full 3D correlation between modalities^4^. Correlative light and electron microscopy (CLEM) approaches have advanced significantly, particularly with cryogenic workflows that preserve native ultrastructure^4-7^. Yet, current cryogenic CLEM (cryoCLEM) techniques face limitations in thick, multicellular, or hybrid tissue samples. Room-temperature 3D CLEM methods often rely on chemical fixation, heavy metal staining, or plastic embedding, all of which compromise the native composition and morphology of the sample^4^. Even in cryogenic workflows, precise 3D targeting across imaging modalities remains limited, particularly when integrating fluorescence and spectroscopic information with electron microscopy (EM)^5^.

To address this, we developed a live-to-cryo correlative imaging workflow that combines live fluorescence microscopy, live and cryogenic Raman microscopy, targeted cryogenic FIB/SEM and cryogenic TEM including electron diffraction, EDX, and tomography. This pipeline allows 3D registration of biological, chemical, and structural information spanning from millimeter-sized tissues to nanometer-scale components without compromising sample integrity.

We demonstrate this method using regenerating zebrafish (ZF) scales as a benchmark system for studying collagen organization and mineralization in the extracellular matrix. This system enables us to showcase the workflow’s ability to resolve collagen fibril orientation, density, as well as mineral crystal shape, orientation and composition at specific 3D locations within a hydrated, unembedded tissue sample. The rigorous 3D correlation of different imaging modalities originating from both life science and materials science provides a new promising strategy for the uncompromised, multiscale analysis of structural biological materials and hybrid biomaterial interfaces.

## Results

### 1. Live imaging of ZF scales

The zebrafish scale has been proposed as model system for studying bone regeneration ^8, 9^. Its greatest advantage is the fish’s ability to generate a new scale when needed, providing a unique opportunity to study the structural and developmental aspects of a newly generated hybrid material tissue. The scale anatomy is mainly composed of a large region of mineralized and non-mineralized collagen. The external (outward facing) mineralized surface is divided into segments by non-mineralized collagen grooves named radii, radiating out from the anterior area that is embedded in the fish tissue, to the posterior edge of the scale (XY view Fig. 1A,B). At the posterior edge, the epidermis cell layer separates the scale from direct contact with the water outside^10^. When viewed in cross-section (XZ view) around the radius (grooves), the scale is organized in 3 compositionally different layers: *i)* a mineralized collagen layer (external layer), *ii)* a non-mineralized collagen layer (elasmoid), and *iii)* a layer of cells that produce and maintain the extracellular matrix (elasmoblasts)^8^. In this work, the cross-sectional plane perpendicular to the direction of the radius will be denoted as the XZ plane, and the cross-sectional plane parallel to the direction of the radius as the YZ plane, regardless of the imaging technique used (Fig. 1A).

**Figure 1.**
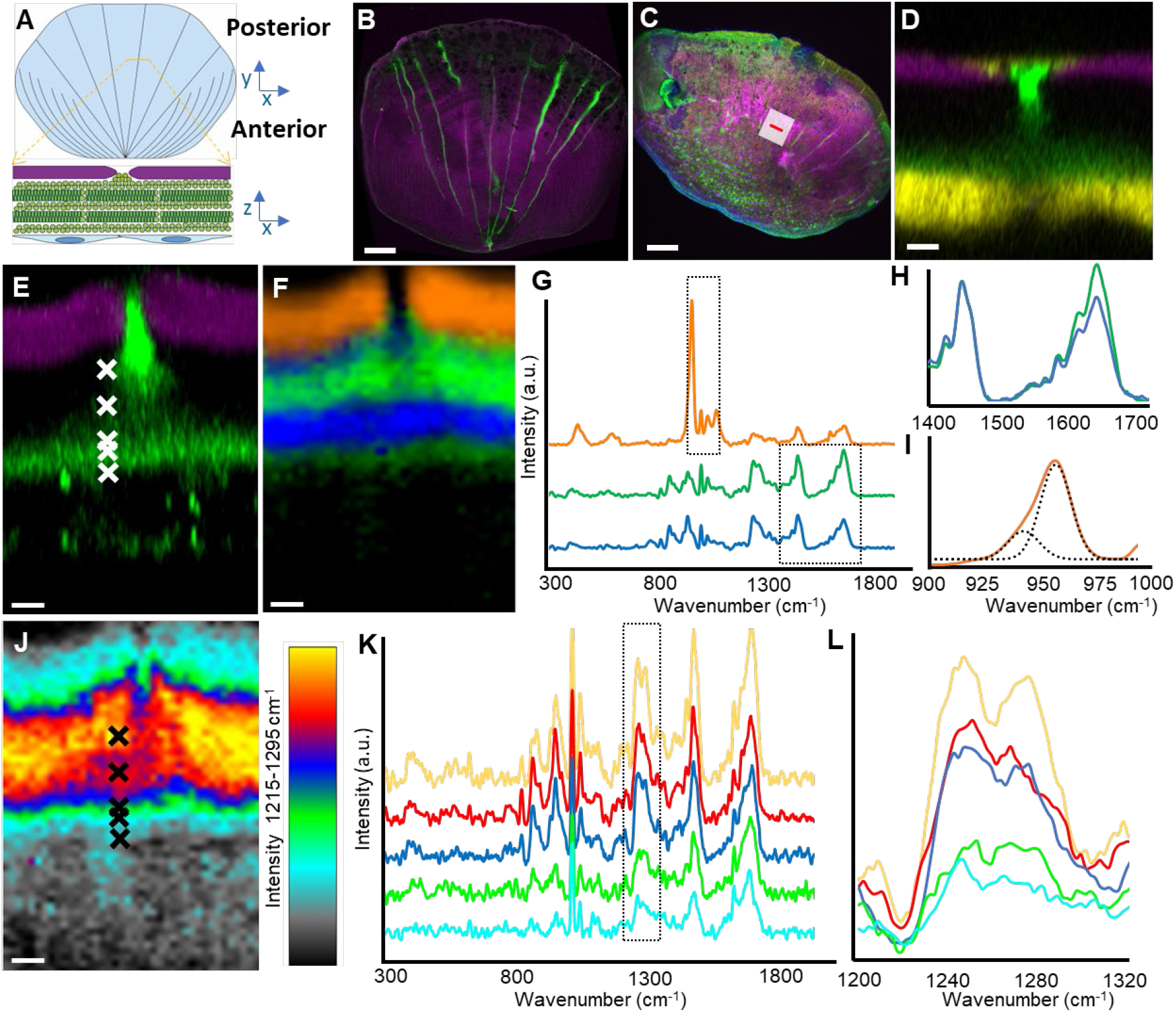
Correlative Live Airyscan confocal microscopy and Raman microscopy of ZF Scale. A) Schematic overview of (top) a full ZF scale in the XY view showing the radii and (bottom) XZ view of a single radius indicating the mineralized external layer (purple), collagenous elasmoid (green), and elasmoblasts (blue). B) Live ACM XY view of a mature fish scale (ZF scale 1). Purple: mineral (calcein), green: collagen (CNA35-mCherry) C-D) Live ACM of a fish scale extracted after 6 days of regeneration (ZF scale 2; Supplementary Fig 1A). Purple, green as in (B), yellow: SP7-expressing cells (eGFP) and blue: nucleus (Draq5). C) XY view, D) XZ view at the location indicated in (C). E-L) ACM-Raman correlative imaging of a fish scale (ZF scale 3; Supplementary Fig. 1B). E) Live ACM XZ view. Colors as in (B). F-I) Polarized Raman Microscopy of ROI in (E). F) Raman component image. Orange: mineralized collagen (external layer), blue and green: elasmoid collagen oriented parallel and perpendicular to the laser, respectively. G) Color-matched Raman spectra of components indicated in (F). H) Intensity of orientation-dependent Amide I vibration (1666 cm^-1^) normalized to the CH_2_ vibration (1455 cm^-1^) indicated in (G), I) Deconvolution of the PO_4_ v_1_ vibration (960 cm^-1^) indicated in (G). J-L) Raman intensity data from ROI in (E). J) Heat map of the matrix density measured using the Amide III vibration (1255 cm^-1^), location matched with (E). K) Color-matched Raman spectra of intensities indicated in (J), recorded at locations (X) indicated in (J) and (E). L) Zoom in on the amide III vibration (1255 cm^-1^). Scale bars: B-C: 200 µm, D: 5 µm, E, F, J: 3 µm.

To investigate the relationship between the structural and compositional build-up of the scale during scale development, we investigated the radii region in scales extracted after 6 days of regeneration (Fig. 1C, Supplementary Fig. 1). The scales were genetically labeled with green fluorescent protein (GFP), indicating active elasmoblasts via SP7 (Osterix) expression ^9^. In addition, chemical live fluorescent staining was used to label the mineral (calcium phosphate - calcein blue) and type I collagen (CNA35-mCherry)(Fig. 1B,C)^11^.

### 2. 3D Correlative imaging connecting fluorescence, Raman, and electron microscopy

In a ZF scale harvested after 6 days of regeneration (ZF scale 2, Fig. 1C, Supplementary Fig. 1A), areas of active matrix formation were targeted by localization of SP7-positive elasmoblast cells around the radii using live 3D airyscan confocal microscopy (ACM). The CNA35-mCherry collagen labeling was selective for the youngest, last formed collagen layer (adjacent to the cell layer) and for the (complete) radii (green in Fig. 1D), implying differences in structure and/or composition between the different collagen layers of the elasmoid.

To further investigate the structural and chemical differences among the collagen layers, we used Raman spectroscopy, which enables label-free imaging of molecules. The correlation of 3D live ACM and live Raman microscopy (2D depth scan) produced accurate overlays of the respective signals for the elasmoblast and mineral layers in the XZ cross-sectional plane (Fig. 1E-L, ZF scale 3, Supplementary Fig. 1B). The collagen orientation in the different layers was visualized using polarized Raman microscopy, exploiting the orientation dependence of the amide I (C=O) vibration (1666 cm^-1^; Fig. 1G,H)^12^. Live Raman depth scan of the elasmoid layer, visualizing the plywood-like structure, shows multiple continuous layers of oriented collagen, rotated by ∼60° relative to one another. The layers cross the radii region without changing orientation. (Fig. 1F).

A Raman heatmap of the orientation *independent* amide III (O=C-NH) vibration (1255 cm^-1^) revealed low collagen signal intensities in both the radii and the youngest collagen layers close to the cell layer that co-localized with the fluorescently labelled collagen, contrasting the more intense Raman signals of the unstained, more mature collagen between the radii (Fig. 1J-L). Live Raman microscopy also visualized the external layer, where the collagen is mineralized with carbonated hydroxyapatite (cHAp, Fig. 1F,G). Here, the collagen density was found to be ∼ **50%** compared to the two neighboring elasmoid layers (Fig. 1K,L). In contrast to earlier reports^11, 13^, the collagen plywood organization, was found to extend also throughout this mineralized layer (Supplementary Fig. 2).

The mineral-to-matrix ratio (M/M) in the external layer - as defined by the relative intensities of the orientation independent PO_4_ v_4_ (432 cm^-1^) mineral and amide III (1255 cm^-1^) collagen vibrations - varied between 0.3 and 1.1, similar to what is found for developing ZF Bone ( Supplementary Fig. 3)^14, 15^. Detailed analysis of the spectra revealed the PO_4_ v_1_ vibration of the cHAp at 960 cm^-1^ had a significant shoulder at 950 cm^-1^ (Fig. 1I) indicating the presence of an acidic calcium phosphate precursor phase, as was previously proposed from energy-dispersive X-ray spectroscopy data ^16^, yet no details are currently available on the structural characteristics of this phase, on its interplay with cHAp, nor on how it is embedded within the collagen matrix.

Although Raman microscopy indicated that matrix density varied across the different matrix regions independently of collagen fibril orientation (Fig. 1F,J), it did not resolve the nanoscale details of the structural organization in these domains.

To obtain detailed insight into the organization of collagen and mineral in the ZF scale, we set out to achieve 3D nanoscale structural information in both the elasmoid and the external layer using EM. The accurate placement of this nanoscale EM information within the macroscopic context of the ZF scale requires a 3D correlative approach with a sample preparation strategy that preserves the scale’s ultrastructure in a near-native state. CryoCLEM, combining 3D cryoACM and 3D cryoFIB/SEM, is a promising approach for deformation-free, high-resolution 3D imaging of tissue samples^5^. However, the 3D targeting accuracy in this approach is limited in the detailed 3D overlay of EM and fluorescence data due to the relatively low z-resolution (∼1000 nm) achievable with cryoACM. As live ACM does not experience the optical aberrations introduced by the air-ice interface in cryo-confocal microscopy^5^, we developed a workflow to directly correlate the 3D cryoFIB/SEM data with the high-resolution 3D information from live ACM (Supplementary Fig. 6, see material and methods for details).

Here we used the FinderTOP pattern in cryo-reflection microscopy for the coarse XY alignment of the sample in the FIB/SEM. Next, we overlay of information from live ACM with the secondary electron (SE) signal of cryoFIB/SEM to navigate to a predefined region of interest (ROI) (Fig. 2A, Supplementary video 1^17^). Subsequently, we use 3D live ACM, to target a groove in the external layer, where the radius spanned the space between an SP7-expressing cell population on the posterior side of the scale and the early-stage collagen above the elasmoblast layer (Fig. 2B, for details see materials and methods). 3D cryoFIB/SEM imaging revealed the structure of the scale, including the external layer, four distinct collagen layers, the radius, and the elasmoblast layer, with nanoscopic detail (voxel size 20x20x40 nm) (Fig. 2C, Supplementary video 2^17^). The combination of aberration-free live ACM and deformation-free cryoFIB/SEM yields a 3D map that enabled direct correlation of information from both modalities at any point within the investigated volume (Fig. 2D) with a average registration accuracy between 63 and 623 nm in the XZ plane (Supplementary Fig. 4) and visualization of the fully correlated 3D volumes (Fig. 2E, Supplementary video 2^17^) within the macroscopic context of the ZF scale.

**Figure 2:**
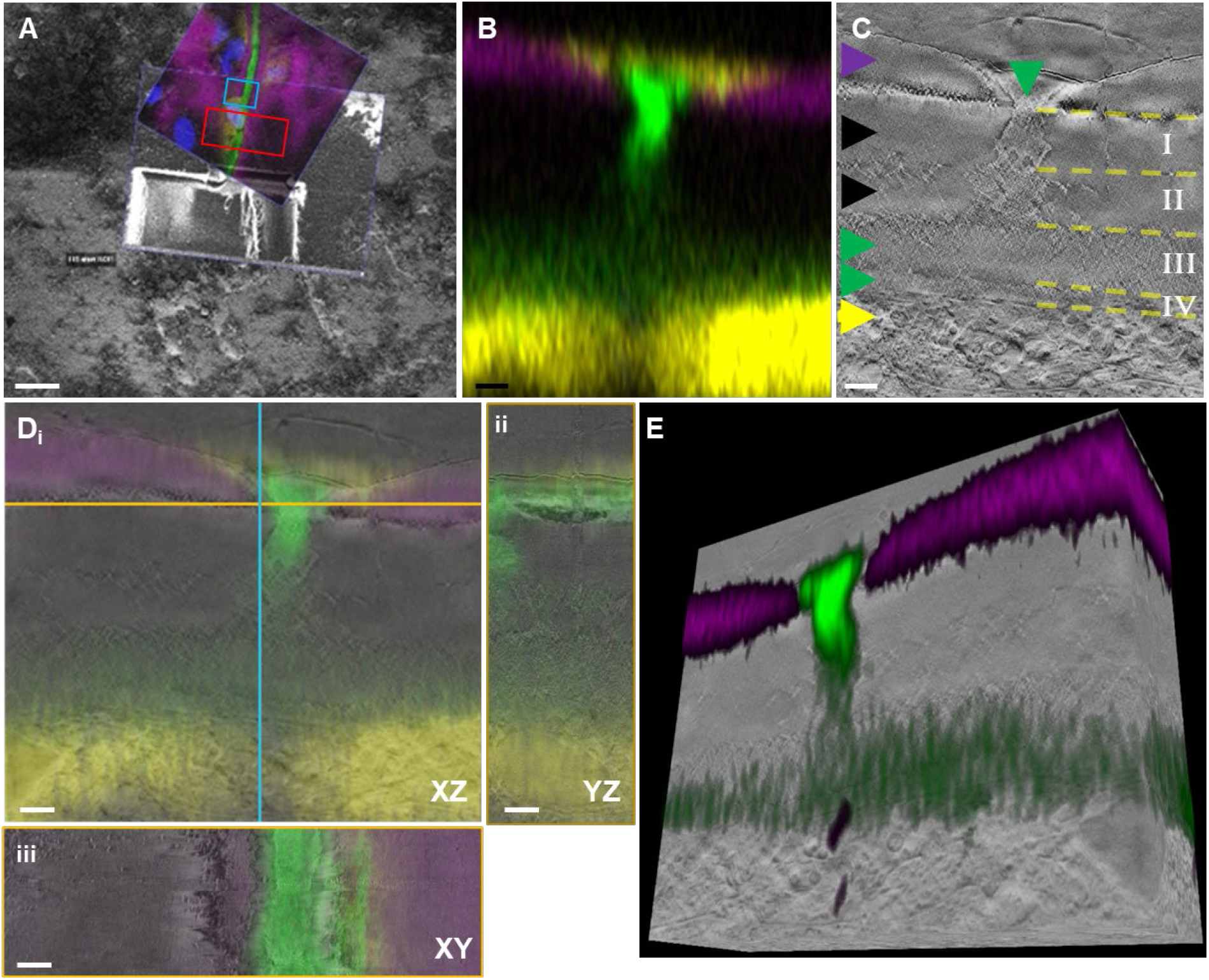
3D Cryogenic correlative light and electron microscopy of a ZF scale. (ZF scale 2, Supplementary Fig. 1A). A) Alignment of cryoSEM and live ACM XY views before targeted cryoFIB/SEM serial imaging. The red box indicates ROI used in panel B-E and Fig. 3A-E. Blue box indicates the location of the ROI used in Fig. 3F-I. B) live ACM XZ view image recorded in the red box in (A). Purple: mineral (calcein), green: collagen (CNA35-mCherry), yellow: SP7-expressing cells (eGFP) C) Secondary electron XZ image from cryoFIB-SEM serial imaging location-matched to the live ACM in (B). Yellow lines indicate borders between collagen layers in elasmoid; Horizontal arrow heads indicate: external layer (purple), unstained elasmoid (black), stained elasmoid (green), elasmoblasts (yellow), vertical green arrow head: radius. D) 3D overlay of live ACM and cryoFIB/SEM stack. i: XZ plane, ii: YZ plane, iii: XY plane. E) 3D overlay of ACM and cryoFIB/SEM volumes, view rotated 180 ^°^ around the z-axis with regard to (B&C). Scale bars: A: 25 µm B-D: 5 µm.

### 3. 3D cryoFIB/SEM reveals collagen fibril organization in the elasmoid layer

To investigate nanoscale structural differences in collagen organization across the different layers, we targeted the area around the radius using 3D cryoFIB/SEM, based on the radius’ strong fluorescent signal (Fig. 2B, Supplementary Fig. 1A). In the cryoFIB/SEM images, the radius and the youngest collagen layer showed a grainy appearance, in contrast to the denser collagen layers outside the radius that showed a low homogeneous contrast (Fig. 3A). As Raman microspectroscopy had demonstrated that the composition of these layers only differed in matrix density (Fig. 1J-L), we attribute these contrast differences to locally varying electron-material interactions related to the relative amounts of tissue fluid present. Within the radius we discerned four collagen layers, which in the XZ plane each presented a distinctly different fine structure (Fig. 3B). However, viewing the same layers in the XY plane (Supplementary video 3^17^, movie through z) showed that their apparently different structures all represented layers of very similar well-aligned fibrils, that only differed in their orientation with respect to the viewing direction (Fig. 3C). Specifically, the layers had angular offsets with the long axis of the radius of 0°, 60°, 150° and 210°, respectively, (Fig 3D-E), and of 60° (*I - II*), 90° (*II - III*) and 60° *(III - IV*) with each other (Fig. 3E).

**Figure 3:**
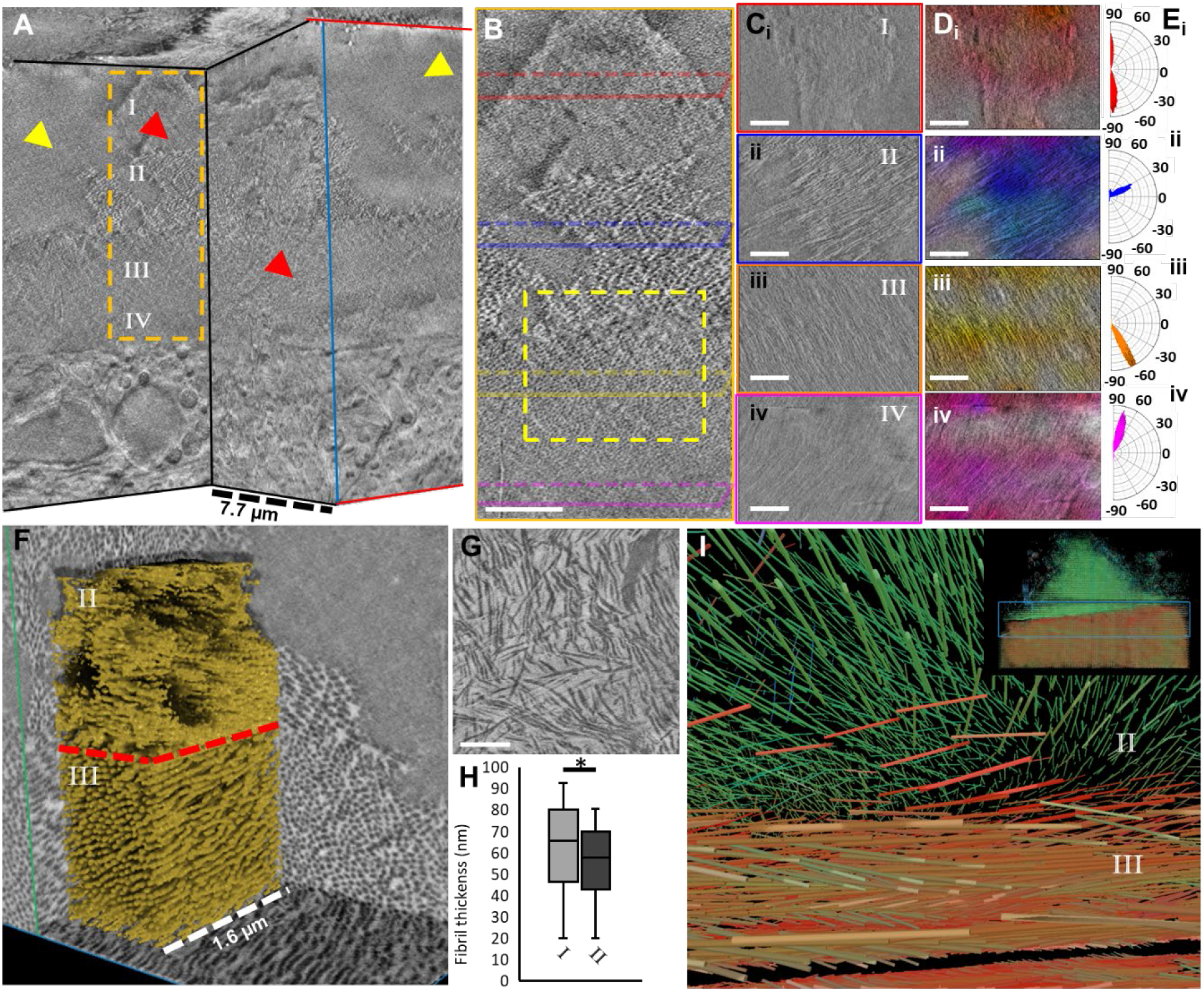
CryoFIB/SEM of collagen organization in the radius of a ZF scale. (ZF scale 2) A) 3D representation of the cryoFIB/SEM volume image shown in Figure 2. Red Arrow indicates the radius and the newest collagen layer (grainy appearance), yellow arrow indicates the denser collagen layers outside the radius (homogeneous contrast). Inset: XZ view of the ROI highlighted in (B). I-IV indicate different elasmoid collagen layers as in Figure 2. B) Zoom in on the area indicated in (A). Colored parallelograms indicate the planes in (C) & (D). C) XY views of cryoFIB/SEM of the planes indicated by the red (i), blue (ii), yellow (iii), and pink (iv) lines in (B). D) Orientation-based false-colored images (Fiji OrientationJ Analysis plug-in) of the different collagen orientations in (C). E) Plots of the direction distributions in (D). F) High-resolution 3D cryoFIB/SEM (voxel size 5x5x5 nm) recorded in the ROI indicated in (B). Inset: Machine learning assisted segmentation and 3D rendering of individual collagen fibrils. The red dashed line indicates the layer II / layer III interface as in (D). G) CryoFIB/SEM image of the interfacial layer indicated in red in (F). H) Collagen fibril diameter in layer II and III based on segmentation in (F). I) Eigen vector analysis (Dragonfly) of segmented collagen fibrils in (F) *p=0.05. Scale bars: B,C and D: 2 µm, G: 1 µm.

To resolve the organization of the collagen fibrils at the level of individual fibrils, we acquired a high resolution 3D cryoFIB/SEM stack (voxel size 5 nm, volume 7.5 x 6.0 x 7.1 µm) at the interface of two adjacent layers in the collagen plywood structure within the volume of the radius (Fig. 3F, Supplementary video 4^17^) The stack was subjected to machine learning-based automatic segmentation (Fig. 3F, inset; Supplementary video 5^17^ ; see materials and methods for details). This segmentation revealed average fibril thicknesses of 55 ± 17 nm (n=56) and 63 ± 19 nm (n=80) for layers II and III, respectively (Fig. 3H and Supplementary Fig. 5). Eigen vector analysis^18^ showed that the individual fibrils were well aligned within the layers and that the adjacent layers had a rotational offset of 90 degrees (Fig. 3I), in line with the lower resolution images (Fig. 3E). Misaligned fibrils were only observed in the single-fibril-thick interfacial layer between the upper and lower layers. In this interfacial layer, the packing density of the collagen fibrils was significantly lower, and the orientation of the fibrils was random (Fig. 3G).

### 4. CryoTEM and CryoET reveal collagen packing gradient in the elasmoid layer

As the low cryoFIB/SEM contrast in the dense, mature collagen areas outside the radii did not allow us to resolve collagen organization at the fibrillar level, we employed cryoTEM, which exploits a different electron imaging contrast mechanism. A cryoTEM lamella was prepared by cryoFIB milling on a pre-defined position in a ZF scale (ZF scale 4, Supplementary Fig. 1C), guided by information from 3D live-ACM and 3D cryoFIB/SEM (Fig. 4A; Supplementary Fig. 6). After lift-out and mounting on a half-moon grid under cryogenic conditions, the lamella was thinned to a thickness of ∼ 200 nm, i.e., thin enough for electron transparency but thick enough to contain a sufficiently large volume for cryogenic electron tomography (cryoET) (Supplementary Fig. 7; Supplementary video 6^17^)

**Figure 4.**
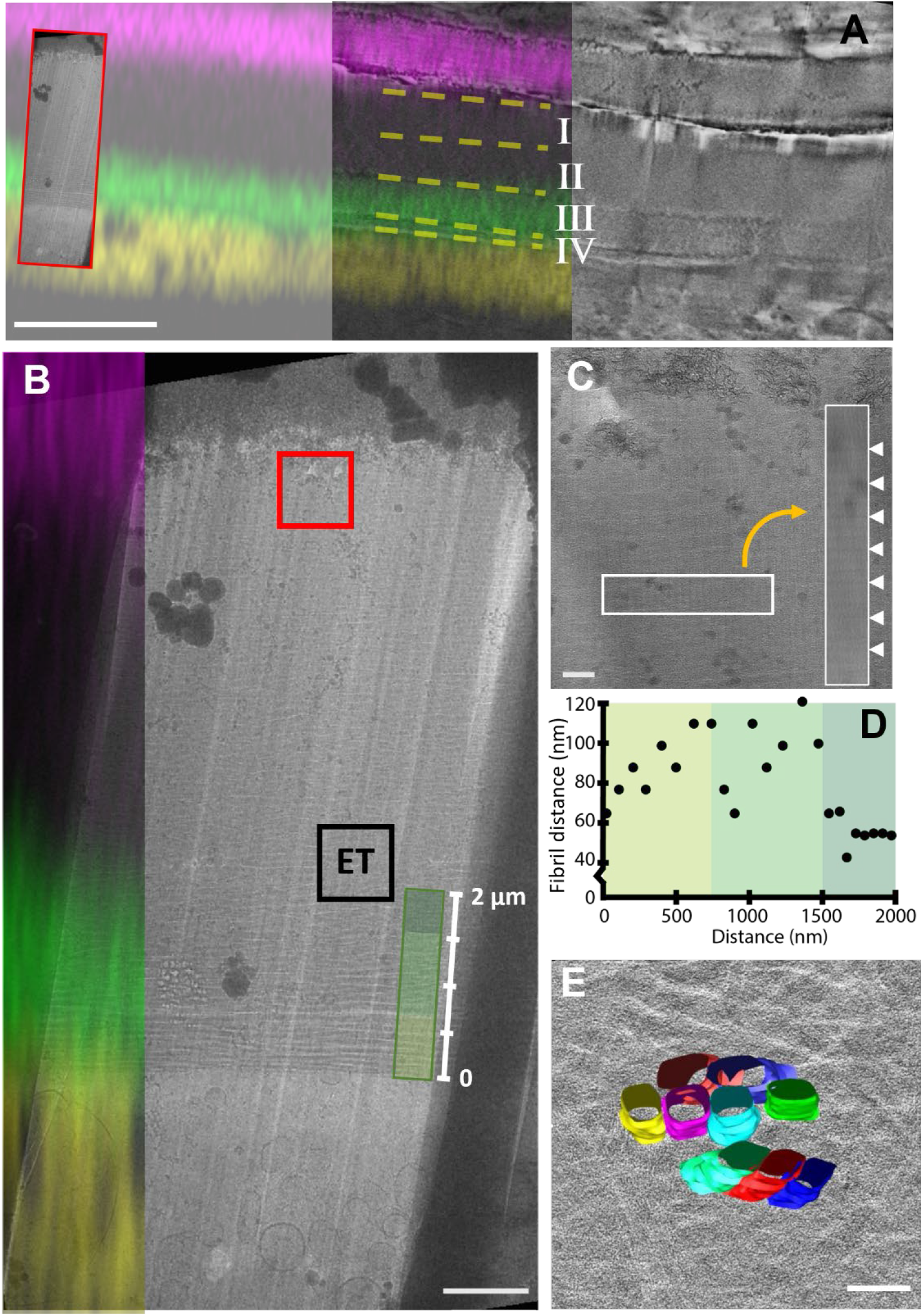
3D Correlation of live ACM, cryoFIB/SEM and cryoET in a ZF scale. (ZF scale 4). A) Location matched XZ plane from live ACM (left and middle) and cryoFIB/SEM (middle and right). Inset: cryoTEM of lamella extracted from the location indicated. I-IV indicate different elasmoid collagen layers as in figures 2 and 3. B) cryoTEM overview (right) of the lamella indicated in (A) location matched with live ACM (left). Red and black boxes indicate areas analyzed in (C) (D) and (E), respectively. C) CryoTEM of the collagen-mineral interface. The arrowheads in the inset indicate a collagen D-band pattern. D) Plot of distance between collagen fibrils starting from the cell-collagen interface. E) 3D reconstruction of the cryo-electron tomogram recorded at the interface between collagen layers II and III (Supplementary Video 7^17^) with segmentation of selected collagen fibrils at the interface. Scale bars: a: 5 µm, b: 1 µm, c: 200 nm, e: 100 nm.

Benefitting from the high Z-resolution in the live ACM data, ROIs were identified in the cryoTEM overview (Fig 4B) at the mineral-collagen and cell-collagen interfaces and at the interface of the fluorescently stained and the unstained collagen layers (Fig. 4B, Supplementary Fig. 8). Higher resolution cryoTEM images of these ROIs resolved the individual collagen fibrils as well as their typical banding pattern (Fig. 4C), and revealed the organization in the different layers (Fig. 4; Supplementary Fig. 8), with thinner, loosely packed collagen at the interface with the elasmoblast layer and a dense packing of thicker fibrils at the interface with the external layer. The fibrils found at the cell-collagen interface had an average thickness of 37 ± 4 nm (n=4), whereas the stained fibrils at the stained/unstained collagen interface had average thicknesses of 67 ± 10 nm (n=4), and the unstained fibrils at the collagen-collagen interface had an average thickness of 86 ± 12 nm (n=6) (Supplementary Fig. 8).

The border between the stained and unstained layers exhibited an interfacial single layer of fibrils (Fig. 4E), similar to what we observed in the radius (Fig. 3G). The orientation of individual collagen fibrils in a 3D volume of 1.0 x 1.0 x 0.2 µm probed by cryoET showed the relative orientations of the stained and unstained main layers to be 60 degrees, and the single intermediate layer to locally have an angular offset of ∼30 degrees with the two main layers in this region (Supplementary video 7^17^).

### 5. Structure and chemistry of the mineral in the external layer

Next, we aimed at using cryoTEM to investigate the detailed structural and morphological characteristics of the acidic phosphate-rich mineral phase in the external layer. To this end, we used the overlay of live ACM (XY view), cryoRaman (XZ view), and cryoFIB/SEM (XZ view) to target an ROI in the external layer of a high-pressure frozen ZF scale (Supplementary Fig. 1D) for the lift-out of a cryoTEM lamella (Fig 5A-C). Both the cryoFIB/SEM back-scattered electron (BSE) images and the cryoRaman depth scan (XZ plane) identified the external layer and the elasmoid by indicating distinct mineralized and non-mineralized collagen layers, respectively (Fig 5C). Spectral deconvolution of the live Raman PO_4_ v_1_ vibration in the mineralized layer, confirmed the presence of acidic phosphate in the external layer microscopy (Fig. 1I), and indicated that the signal associated with acidic phosphate accounted for ∼ 35% of the total PO_4_ v_1_ peak intensity (Fig. 5 E,F) (for detailed analysis of Raman data see Supplementary Fig. 9).

**Figure 5.**
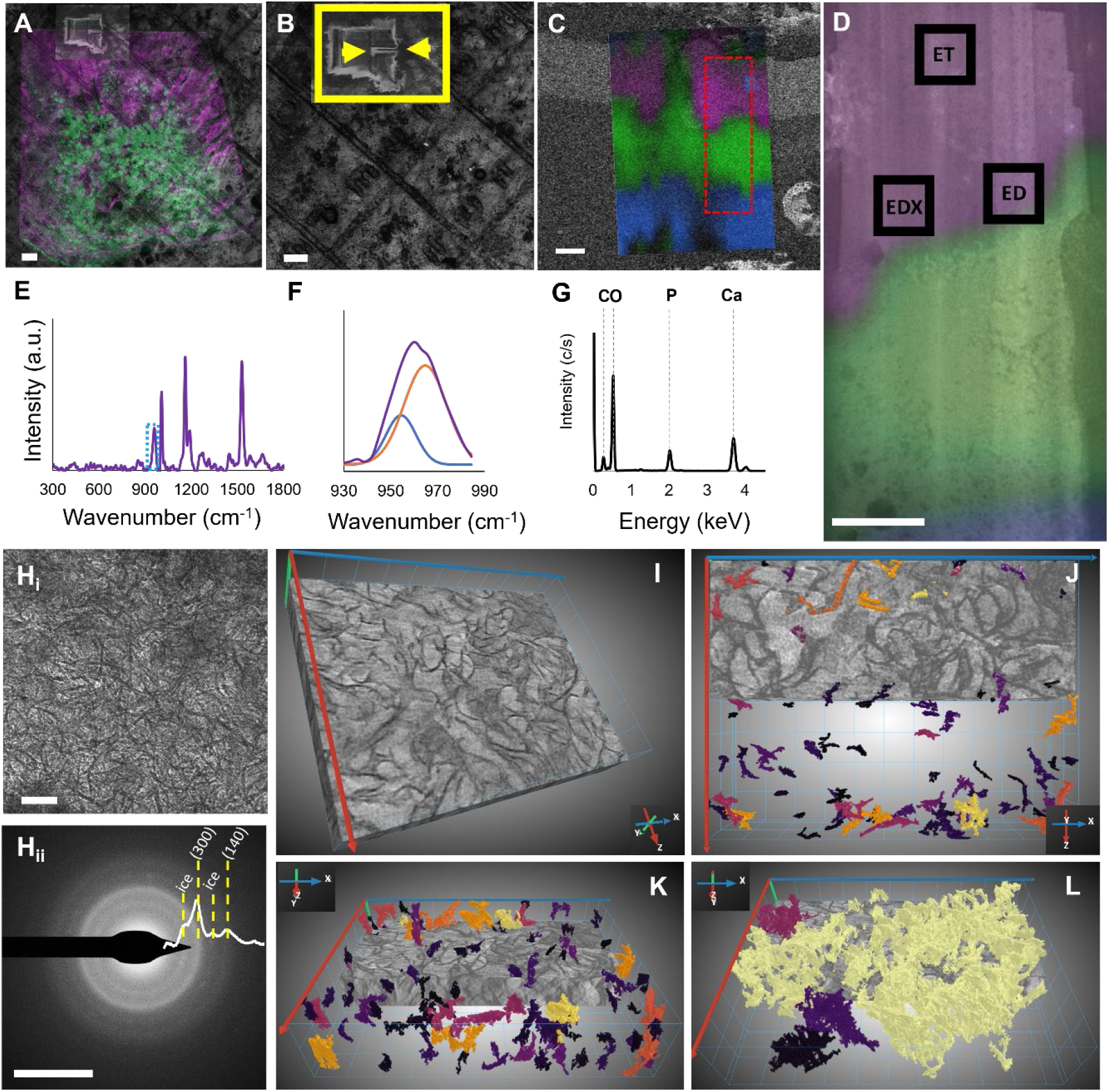
3D Correlation of live ACM, cryoRaman, cryoFIB/SEM and cryoET in a ZF scale. A) Overlay of live ACM and cryoFIB/SEM XY-view showing the FinderTop and trench for lift-out. B) Zoom-in of (A) on lamella. Yellow arrows indicate the position of the cryoRaman depth scan. C) Position matched overlay of polarized cryoRaman micrograph of depth scan and cryoFIB/SEM BSE image of the location of the lamella prior to lift out. The red box indicates the lift-out window used for cryoTEM. Raman spectral components indicate an external layer (purple) and elasmoid layers with different orientations (green, blue). D) Overlay of cryoTEM on lift-out and cryoRaman depth scan from (C). Boxes indicate areas where tomogram (ET), energy dispersive x-ray (EDX), and selected area electron diffraction (ED) were collected. E) Average cryoRaman spectrum of external layer. F) Zoom in on the box in (E) with the deconvoluted PO_4_ v_1_ peak. G) EDX spectrum on location indicated (D). H) CryoTEM (i) and SAED (ii) of the location indicated in (D). I) Reconstruction of the tomogram recorded at the location indicated. Individual mineral particles are segmented and indicated in different colors. (D). J) XZ view of the reconstructed tomogram showing particles with dimensions < 100 nm. K) Reconstructed tomogram tilted in the Y direction with respect to (L). L) Reconstructed tomogram with segmentation of the four biggest continuous mineral networks. Viewing direction as in (K). Scale bars: A, B) 50 µm, C, D) 2 µm, H_i_) 50 nm, and H_ii_) 5 nm^-1^

Overlay of the cryo-Raman depth scan and the cryoTEM overview of the lamella after thinning identified the external layer and two neighboring elasmoid collagen layers (Fig 5D). This allowed targeted recording of higher resolution 2D cryoTEM images (Fig 5H_i_), tomograms (cryoET, Fig. 5I-L), low dose selected area electron diffraction (LDSAED, Fig. 5H_ii_) patterns, and cryo-electron dispersive X-ray (EDS) spectra (Fig 5G) in the external layer, collecting nanoscale morphological, structural, and chemical information on the mineral in precisely defined areas where also Raman spectroscopic information was obtained.

Electron tomograms revealed that the mineralized external layer contained a 3D network of intergrown curved plate-like crystals (Fig. 5I-L, Supplementary video 8^17^). When viewing the tomograms perpendicular to the XZ planes of the ZF scales (XY view in the tomogram), the crystals most frequently appeared face-on, whereas they did not when observed along the XY or YZ planes. In line with these observations, LDSAED patterns recorded in the XZ plane of the scale (XY plane in cryoTEM) showed diffraction patterns that were consistent with cHAp but that were lacking the characteristic (002) reflection, indicating a preferred orientation of the cHAp c-axis near-parallel to the electron beam, hence within the plane of the external layer (Fig 5H)^19^. These observations are consistent with the lacey patterns reported for bone samples when imaged parallel to the long axis of the mineralized collagen fibrils^20^.

CryoEDX spectra displayed peaks for Ca, P, O and C, with a Ca/P ratio of 1.60-1.62 (Fig 5G), which is low compared to cHAp (Ca/P = 1.83-1.97; CO_3_ substitution 4-8 mol%) but completely in line with an average overall molecular formula Ca_9.24_(PO_4_)_2.82_(CO_3_)_1.52_(OH)_1.45_(HPO_4_)_2.77_ (Supplementary Fig. 10) that represents cHAP with 27-33 mol% incorporated acidic phosphate.

## Discussion

We developed and applied a live-to-cryo correlative imaging workflow that combined multiple imaging and analytical modalities: live fluorescence microscopy, live and cryo Raman microscopy, cryoFIB/SEM, cryoTEM, cryoET, cryoEDX, and LDSAED. We demonstrate the correlation among these microscopy techniques and show how they can be used to obtain qualitative and quantitative information on the chemistry and structure of mineralizing tissues at predefined points within their 3D hierarchical structure. This approach enables imaging across multiple length scales, zooming in over four orders of magnitude - from observing the millimeter dimensions of the entire tissue in light microscopy to the nanometer-scale structures in cryoTEM - while collecting biochemical, chemical, and crystallographic information with ACM, Raman microscopy, EDX, and electron diffraction.

The zebrafish scale, which has been proposed as a model system for studying bone formation ^21, 22^, was selected as a demonstrator system to benchmark the live-to-cryo correlative workflow. Although it has been studied for decades, previous reports have presented limited details and, in some cases, inconsistent findings regarding the chemistry and structure of both the mineralized and non-mineralized parts of the matrix ^11, 13, 18, 23-28^. As these discrepancies are most likely due to differences in methodology and sample preparation, we emphasize that the results presented here were in all cases obtained from zebrafish scales extracted after 6 days of regeneration, following non-destructive sample preparation.

The plywood structure of the elasmoid (Fig. 3) was already revealed in earlier work^13, 24^, but due to limited technical possibilities at that time, particularly due to the sample preparation required for EM, details on the matrix density and the organization in the radii had not been revealed. The selective staining with CNA35-mCherry of type I collagen indicate there are differences in the structure or composition of collagen in the layer closest to the cell and radii compared to outside the radii. Raman microscopy revealed differences in collagen density between the stained and unstained regions. This was further confirmed by cryoFIB/SEM that showed the radii consisted of loosely packed collagen fibrils, with diameters and cross-sectional densities that varied across layers. Additionally, cryoTEM showed that outside the radii, the fibril diameters and cross-sectional densities increased stepwise from layer to layer. Raman microscopy revealed that the density variations between adjacent layers were independent of their angular offset, which persisted across the cross-sectional boundary of the radius.

Previous literature also presents a scattered view of the structure and chemistry of the mineral in the external layer of elasmoid scales. Most studies refer to the mineral as hydroxyapatite, but acidic or calcium-deficient precursors have also been found in regenerating scales^26^. The mineral particles have been described as either needles ^11^ or platelets (laths)^28^, and have been reported to be both aligned with collagen orientation ^27^ and randomly oriented ^26^. We attribute these differences, in part, to variations in developmental stage, viewing direction in TEM, and sample-preparation procedures. Using our live-to-cryo multimodal workflow, we unequivocally show that, after 6 days of regeneration, the mineral phase in the external layer of zebrafish scales comprises strongly curved, partially interconnected crystalline platelets composed of a mixed phase of carbonated hydroxyapatite and acidic calcium phosphate. These platelets are oriented parallel to the scale surface and aligned with the collagen fibrils similar as observed in bone^29^.

While individual techniques such as cryoFIB/SEM^4, 30-34^ or cryoET^35-37^ have advanced our ability to image ultrastructure in vitrified samples, they typically lack integrated chemical specificity or require prior identification of target regions in 2D^38^. Correlative workflows often rely on plastic embedding or heavy-metal staining, which can distort morphology and chemistry, particularly in hybrid materials such as mineralizing tissues^3^. By combining live fluorescence and Raman imaging with precise cryogenic targeting and high-resolution electron microscopy, our workflow offers a fully integrated platform for label-assisted 3D targeting, chemical mapping, and structural imaging. This multiscale, multimodal approach enables high-confidence correlation between cellular activity, matrix structure, and mineral composition in intact, hydrated tissues—capabilities not currently achievable with existing single-modality or conventional CLEM techniques.

## Conclusion

The live-to-cryo workflow described here enables determination of the thickness, packing, and orientation of each individual collagen fibril in the elasmoid and each individual mineral platelet in the external layer, thereby providing access to a new level of detail in the organization and composition of the EMC in 3D tissue samples. Beyond its application to zebrafish scales, this workflow is broadly applicable to other hybrid tissues and biomaterials that require correlative, multiscale analysis. Its compatibility with vitrified, hydrated samples enables structural and chemical imaging without embedding or staining, which is essential for preserving native architecture in soft/hard interfaces. The method is particularly suited to studying environmental biomineralization systems (e.g., coral, shells), pathological mineralization (e.g., atherosclerosis, breast tissue calcification), and biomaterial integration (e.g., implants, tissue scaffolds). The modular nature of the workflow, makes it adaptable to diverse tissue types and experimental questions, providing a new platform for structure-function analysis in complex biological systems.

## Acknowledgements

This project was supported by the European Research Council (ERC) Advanced Investigator grants (H2020-ERC-2017-ADV-788982-COLMIN) and (H2025-ERC-2023-ADV-101141998 - REVALVE) and SPARQ grant (Thermo Fisher Scientific) to N.S. A.A. was partly supported by a VENI grant from the Netherlands Scientific Organization NWO (VI.Veni.192.094). EMS was supported by the Ramón y Cajal Program (RYC2023-045512-I) funded by MCIN/AEI/10.13039/501100011033 and FSE+ and the project PID2022-141993NA-I00 funded by MICIU/AEI/10.13039/501100011033 and ERDF/UE.

## Author contributions

Conceptualization: RvdM, LR, MdB, AA, NS

Methodology: RvdM, LR, MdB, RR, DD, JS, AW, BJ, MV, JM, AA, NS

Investigation: RvdM, LR, MdB, RR, DD, JS, AW, BJ, MV

Visualization: RvdM, LR, MdB, DD, JS, AW, AA, NS

Funding acquisition: EMS, AA, NS

Project administration: NS

Supervision: EMS, AA, NS

Writing – original draft: RvdM, LR, MdB, AA, NS

Writing – review & editing: RvdM, LR, RR, JS, AW, EMS, AA, NS

## Competing Interests

The authors declare no competing interests.

## Data availability

Supplementary movies are available in the Zenodo repository (DOI: 10.5281/zenodo.18846011).

## Materials and Methods

### Zebrafish and scale harvesting

Zebrafish were maintained under normal husbandry conditions. Transgenic line Tg(Ola.sp7:NLS-GFP) (DeLaurier, 2010) was kept in an AB/TL background. Anaesthetized fish (0.05% (v/v) tricaine methane sulfonate (MS-222) (UoB), 0.1% (v/v) 2-phenoxyethanol (RU)) were put on a wet tissue containing system water and anesthetic, and scales plucked under a microscope with a watchmaker’s tweezers from the midline of the lateral flanks near the dorsal fin.

### Fluorescent staining

Fresh harvested scales were fluorescently stained to label the mineral (0.1mg/ml Calcein Blue, Sigma, M1255-1G), collagen (2 µM CNA-mCherry^39^) and nucleus (2.5 µM Draq5, Thermofisher, 62251), in the same medium in which the fish were grown (normal husbandry conditions) for at least 2 hours at room temperature. Scales were then rinsed in fish culture medium.

### Fluorescent live-cell imaging

Fluorescent images were recorded using the confocal laser scanning microscope LSM900 (Zeiss microscopy GmbH) with Airyscan 2 Multiplex SR-4Y. Prior to imaging, a Zeiss ZEN Connect project (Zeiss software for correlative microscopy, version 3.5) was created to make a working sheet (canvas) to align and overlay all the images and to facilitate further correlation with other imaging modalities. Images were recorded using the settings according Table 1. First, low resolution images (pixel size 390 nm) were taken for orientation purposes (C Epiplan-Apochromat 5x/0.2 DIC, Zeiss) by creating an image of the entire scale. Next, high resolution 3D images (50x260 nm) were taken of areas surrounding the grooves using a water-dipping lens (W Plan-Apochromat 63/1.0 M27, Zeiss). The scales are vitrified direct after imaging.

**Table 1:**
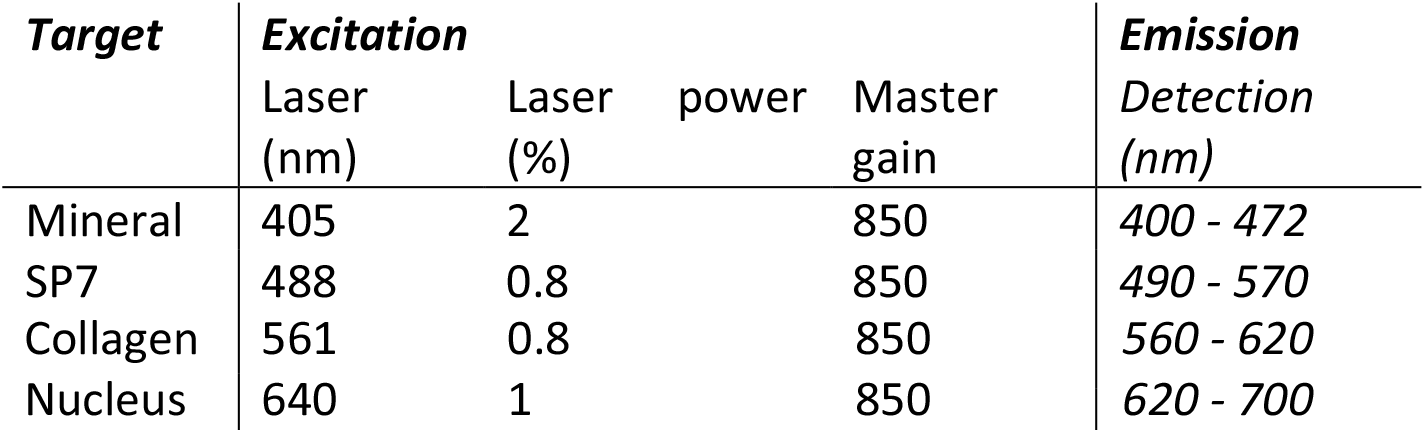
Image acquisition settings.

### Live Raman Imaging

Raman microscopy was performed before (6 days sample) or after (9 and 13 days samples) fluorescent staining/imaging. Staining of the 9 and 13 day scales with the CNA35-mCherry was omitted because of the emission overlap with the Raman excitation at 532 nm.

Samples were measured directly in a 6 wells plate, filled with the same medium in which the fish were grown (normal husbandry conditions). An overview image of the fish scale was first obtained by stitching images from a 10x (Zeiss EC “Epiplan-Neofluar”) objective attached to the Witec Alpha 300R Raman microscope. These images were used for correlation with the reflection and fluorescent images obtained earlier to identify the region of interest. Subsequently, a 63x (W Plan-Apochromat 63/1.0 M27, Zeiss) water dipping objective was used to obtain a detailed image of the region of interest and allow for precise correlation between the two microscopes.

Raman spectroscopy was performed using the same 63x objective as described for live fluorescence at image resolution of 500 nm across a cross sectional area of 25x25 µm. Laser power set at 20 mW for an exposure of 2 seconds per pixel. When CNA35-mCherry staining is used, Raman imaging should be performed prior to the staining and fluorescence imaging to avoid interference of the fluorescent labels with the Raman signals (see SI-1 for details).

### Raman data analysis

The obtained data was analyzed using the project 5 analysis software provided by the supplier (Witec, Ülm). Baseline correction was performed using a rolling ball correction set at radius 300. First unsupervised component analysis was performed to identify the different layers in the ECM as well as the cells and the external layer. Visualization of the component images was done by rounding up the highest identified value of each component to 2 significant numbers, and using this as the maximum value for the image. For the minimum value we used either 10% (collagen and elasmoid) or 30% (cells).

For quantitative analysis the ranges as described in Table 2 were used to determine the orientation (Collagen/Organic matrix), density (Organic matrix) and mineralization (Mineral/Organic matrix) levels of the different regions. Determination of the mineral phase was done by the center of mass of the phosphate v_1_ peak (960 cm^-1^ centre, 60 cm^-1^ width) after subtraction of the collagen signal based on the intensity of the organic matrix peak.

**Table 2.** Raman data analysis settings

| ***Material*** | ***Centre ($\text{cm}^{-1}$)*** | ***Width ($\text{cm}^{-1}$)*** |
| --- | --- | --- |
| Mineral | 590 | 80 |
| Organic matrix | 1455 | 80 |
| Collagen | 1660 | 120 |

### High pressure freezing

Scales were immersed in 10% dextran (31349, Sigma) in water and sandwiched between HPF carriers with 2 mm internal diameter. Prior to freezing all carriers were rinsed in pure ethanol.

For freezing the 0.05 mm cavity carrier (Art. 390, Wohlwend) and a tailor-made grid labeled, flat-sided finderTOP (Alu-platelet labelled, 0.3 mm, Art.1644 Wohlwend) were used, to allow an imprint of a finder matrix on the amorphous ice^40^. The flat side of the finderTOP was treated with 1% L-α-phosphatidylcholine (61755, Sigma) in ethanol (1.00983.1000, Supelco). The samples were then high pressure frozen using live μ instrument (CryoCapCell) and stored in liquid nitrogen.

The HPF carrier containing the vitrified scales were loaded into a universal cryoholder (Art. 349559-8100-020, Zeiss cryo accessory kit) using the ZEISS Correlative Cryo Workflow solution, which fit into the PrepDek® (PP3010Z, Quorum technologies, Laughton, UK).

### Fluorescent cryoimaging

The universal cryoholder containing the samples was transferred into the Linkam adapter to fit the LSM cryostage (CMS-196, Linkam scientific inc.), which was applied to an upright confocal laser scanning microscope (LSM 900, Zeiss microscopy GmbH) equipped with an Airyscan 2 detector. First, overview images with low resolution (pixel size 623 nm) (C Epiplan-Apochromat 5x/0.2 DIC, Zeiss) were made, to visualize the scales with fluorescence microscopy (Table 1) and the ice surface with reflection microscopy. Next, an overlay with the live-cell imaging data was made and the ROIs were recorded in 3D with high resolution (66x360 nm) (C Epiplan-Neofluar 100x/0.75 DIC) including the reflection microscopy images to measure the dept of the scales within the carrier. Settings had to be adjusted for this objective (Table 3).

**Table 3:** Cryolight 100x image acquisition settings

| **Target** | **Excitation** |  |  | **Emission Detection (nm)** |
| --- | --- | --- | --- | --- |
|  | Laser (nm) | Laserpower (%) | Master gain |  |
| Mineral | 405 | 14 | 900 | 400 - 472 |
| SP7 | 488 | 14 | 900 | 490 - 570 |
| Collagen | 561 | 14 | 900 | 560 - 620 |
| Nucleus | 640 | 14 | 900 | 620 – 700 |
| Reflection | 640 | 0.2 | 600 | 400 - 700 |

### CryoFIB/SEM

The universal cryoholder was transferred into the SEM holder in the PrepDek® (Quorum). The sample was sputter-coated with platinum, 5 mA current for 30 seconds, using the prep stage sputter coater (PP3010, Quorum technologies, Laughton, England) and was transferred into the Zeiss Crossbeam 550 FIB/SEM (Carl Zeiss Microscopy GmbH, Oberkochen, Germany) using the PP3010T preparation chamber (Quorum, Laughton, England). Throughout imaging, the samples were kept at -160 °C and the system vacuum pressure was 1 x 10^−6^ mbar.

After inserting the sample into the FIB/SEM chamber, overview images were taken using the SEM to align the data with the LSM reflection image of the surface of the same ZEN Connect project. This alignment enables the stage registration which allows using the fluorescence signal to navigate to different regions of interest. After initial alignment using the SEM, a FIB image of the surface was collected with the 30kV@50pA probe in 54° tilt.

A coarse trench was milled for SEM observation using the 30 kV@30 nA FIB probe. Cold deposition was done with platinum for 30 sec. Fine FIB milling on the cross section was done using the 30kV@700pA probe. For serial FIB milling and SEM imaging the slice (trench) width was 40 μm and for FIB milling the 30 kV@300pA probe was used, with a slice thickness of 20 nm. When a new slice surface was exposed by FIB milling, an InLens secondary and EsB image were simultaneously collected at 2.33 kV acceleration potential with 250 pA probe current. The EsB grid was set to -928 V. The image size was set to 2048 × 1536 pixels. For noise reduction line average with a line average count N = 46 at scan speed 1 was used. The voxel size of the stacks was 20×20×40 (low resolution) or 5x5x5 (high resolution) nm.

### FIB/SEM Data analysis

The cryoFIB/SEM images were processed using MATLAB (R2018b, Natick, Massachusetts: The MathWorks Inc.) to correct for defects such as curtaining, misalignment and local charging. The same software was used for subsequent noise reduction and contrast enhancement. A summary of each processing step is as follows:

#### Curtaining

Removing the vertical stripes in the stacks was done following a wavelet-FFT filtering approach described by^41^. In brief, the high frequency information corresponding to the vertical stripes was successively condensed into a single coefficient map using decomposition by “coif” wavelet family. Subsequently, a 2D-fourier transform was performed to further tighten the stripe information into narrow bands. Finally, the condensed stipe information was eliminated by multiplication with a gaussian damping function and the destriped image was reconstructed by inverse wavelet transform.

#### Alignment

The consecutive slices were aligned using normalized cross correlation. Briefly, the first image in the stack was chosen as reference and the second image was translated pixel by pixel across the reference and a normalized cross correlation matrix was obtained using the “normxcorr2” function. The location of the highest peak in the cross-correlation matrix (representing the best correlation) was then used to calculate the translation required to align the two images. Once the moving image was aligned with the reference image, it served as the reference for alignment of the subsequent slice.

#### Charging

Elimination of the local charge imbalance was achieved using anisotropic gaussian background subtraction. Briefly, the “imgaussfilt” function was used to perform 2D-gaussian smoothing with a two-element standard deviation vector. The elements in the vector were chosen in a manner to apply a broad and sharp gaussian in the horizontal and vertical directions, respectively. Subsequently, the corrected image was obtained by subtracting the filtered image from the original image.

#### Noise Reduction

In order to improve the signal-to-noise ratio, noise reduction was performed using anisotropic diffusion filtering ^42^. Briefly, using the “imdiffuseest” function, the optimal gradient threshold and number of iterations required to filter each image was estimated. Subsequently, the “imdiffusefilt” function was applied with the estimated optimal parameter values to denoise each image.

#### Contrast enhancement

As the final processing step, the contrast was enhanced using “Contrast-limited adaptive histogram equalization”^43^. Using the “adapthisteq” function, the contrast was enhanced in two steps, using a uniform distribution and a low clipping limit in order to avoid over-amplification of homogeneous regions.

#### 3D segmentation

DragonflyTM image analysis and machine-learning software (version 2022.1, Objects Research Systems, Montreal, QC, Canada) was used to segment all image data. 5x5x5 nm images were processed as described above and imported into Dragonfly. 4 slices of the XY view were manually segmented for collagen and background. The embedded Random Forrest machine learning approach was applied to automatically segment the rest of the stack. After segmentation the results were optimized using the opening and smoothing commands to visualize individual fibrils. Thickness mesh was generated on the segmented data with a further lapacian smoothing set at 5, with a thickness range of 25-85 nm. In order to visualize the orientation of the scales, first an anisotropy map was generated in Dragonfly using a threshold of 60 and sampling 2x2x2 voxels. After which a vector field could be generated.

### TEM-lamella preparations

Coarse trenches were milled around the lamella to create access to the thick lamella, using the 30 kV@30 nA FIB probe. After this the lamella was removed from the scale by grabbing it with a gripper (Kleindiek, Reutlingen, nanomanipulator) and attached to a half-moon copper grid (Copper lift-out grids 75964-01, Aurion, The Netherlands ). After the lamella was attached, thinning of the windows was started with using an 30kV@3nA FIB probe, followed by thinning with the 30 kV@700pA and the last polish step was done with the 30 kV@50pA FIB current to make a smooth surface on both sides of the lamella for cryoTEM. Once the windows were approximately 200 nm thick, the grid was transferred to the TEM.

### CryoTEM

Image and tomograms on the collagen were acquired with a Titan Krios present at the Pasteur Institute (courtesy of Matthijn Vos, Paris, France) (Thermo Fisher Scientific, Eindhoven, The Netherlands) equipped with a Falcon direct electron detector (Thermo Fisher Scientific, Eindhoven, The Netherlands) and field emission gun operated at 200 kV. Images and the tomogram on the mineral was acquired with a Talos F200C G2 (Thermo Fisher Scientific, Eindhoven, The Netherlands) equipped with a Falcon 4i direct electron detector (Thermo Fisher Scientific, Eindhoven, The Netherlands) and field emission gun operated at 200 kV. Tomograms were reconstructed and segmented using the IMOD (University of Colorado, CO, USA). Electron diffraction pattern were acquired on a Ceta-S camera (Thermo Fisher Scientific, Eindhoven, The Netherlands) and EDX spectra were acquired with a Bruker X-flash 6T_30 (Bruker, Germany)

## Notes

https://doi.org/10.5281/zenodo.18846011

### Competing Interest Statement

The authors have declared no competing interest.

