## Supplementary material for "Uncompromised, multimodal, multiscale structural analysis of the hierarchically organization in mineralized tissues"

**Supplementary figure 1: Overview of all the used scales in the project**

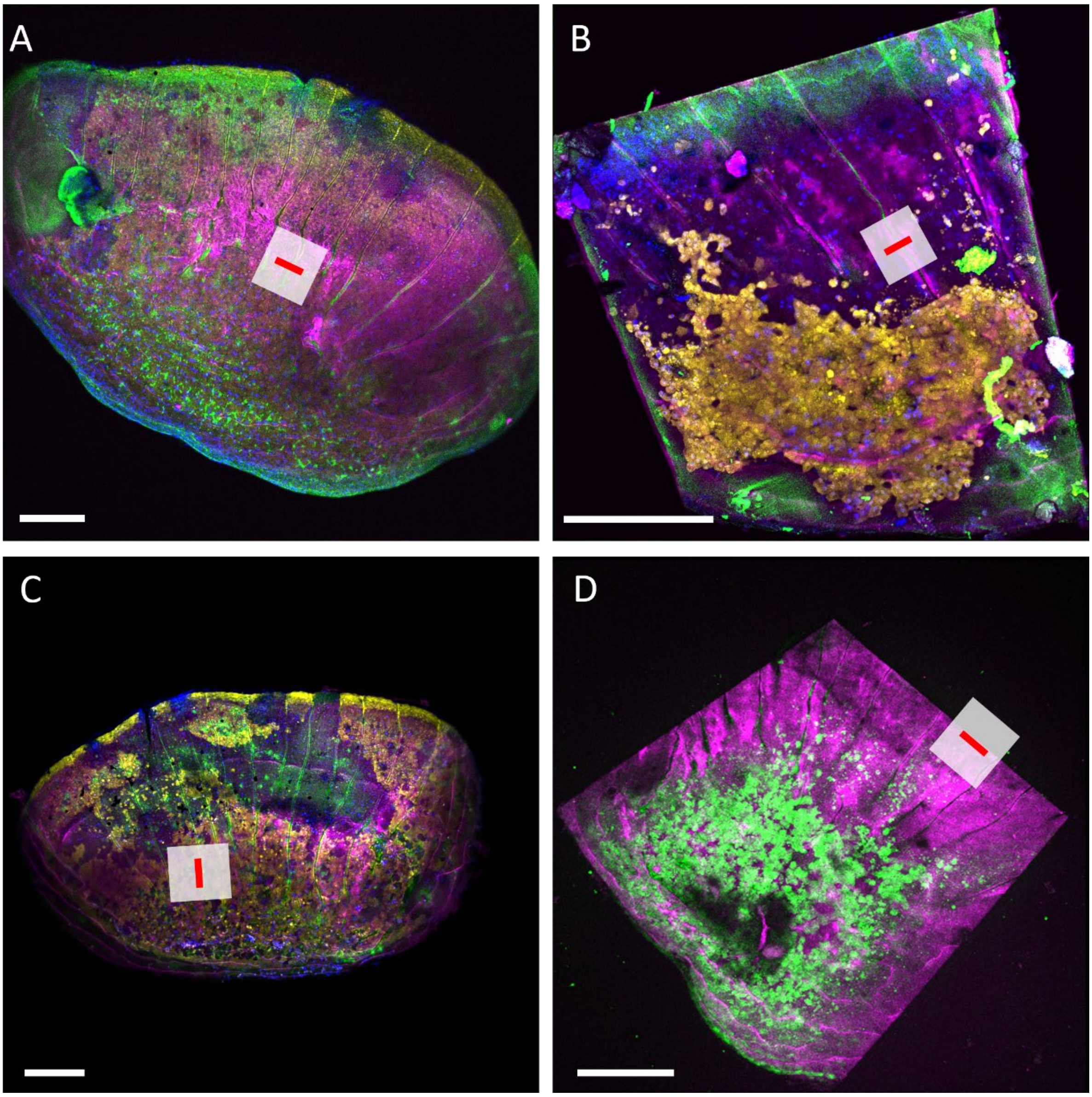


***Supplementary Figure 1*** *Overview of the fish scales in the paper. In each scale the position of the volume imaging and lamella preparation is indicated. Fish scale corresponding to A) figure 1C,D, 2 and 3. B) figure 1 E-F. C) figure 4. D) figure 5. Scale bar 200 µm*

**Supplementary figure 2: Orientation of the mineral layer using Raman microscopy.**

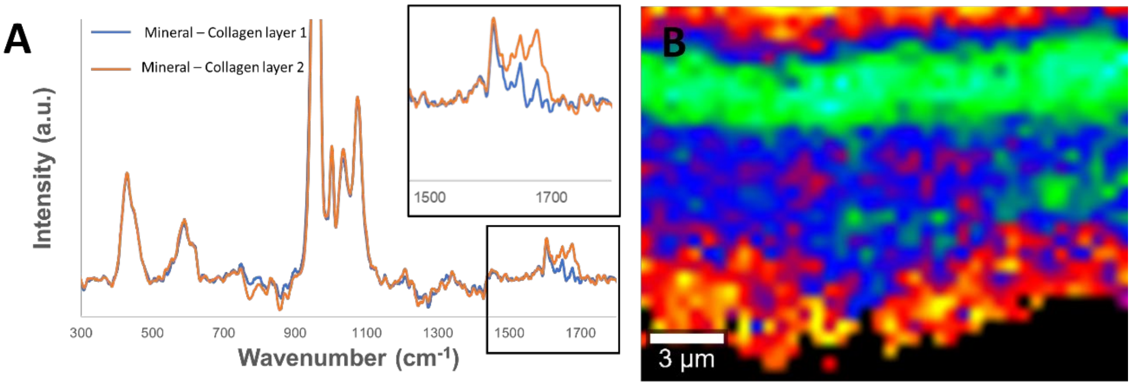


**Supplementary figure 2:** A) Raman difference spectra of the mineralized collagen layer and two unmineralized collagen layers, these spectra show a difference in the height of the polarization dependent amide I peak, indicating a structured organization of the mineralized collagen layer. B) Image of the intensity of the Amide I / Amide III ratio proving different orientations of the collagen layers including the mineralized collagen layer.

**Supplementary figure 3: Mineral density Raman**

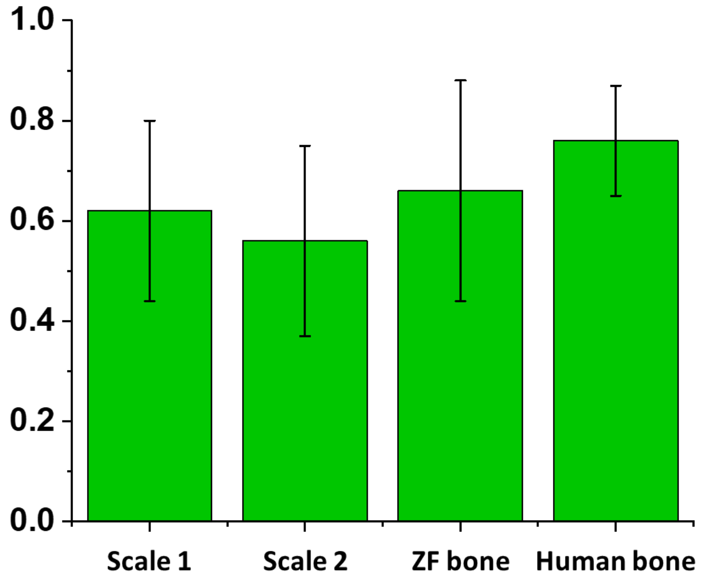


**Supplementary Figure 3:** Mineral-to-matrix ratios, as determined by the ratio of the amide III (1255 $cm^{-1}$) collagen and $PO_4$ $v_4$ (432 $cm^{-1}$) mineral peaks, of two live-measured ZF scale minerals compared to the mineralization of ZF-bone [1] and human bone [2], showing comparable levels of mineralization for ZF scale and ZF bone.

**Supplementary figure 4: Target registration errors**

The target registration error (TRE) was computed for the correlated images with Icy 2.5.4.0 software (Institut Pasteur(Fr)) using the ec-CLEM v.1.1.0.0 plugin.[1] For the EM images, the mineral phase was manually segmented in 3–6 stacks of five consecutive images (mini stacks) in XY, XZ, and YZ views (Supplementary figure 4A). For LSM images, segmentation was done with FIJI (ImageJ v.1.53t) using the automatic threshold function (Supplementary figure 4B). Ten landmarks for registration were chosen on each EM mini stack which were transformed to the corresponding correlated LSM images with ec-CLEM 3D tool (Supplementary figure 4C). For each mini stack, a predicted average registration error map was computed and displayed as a heat map representing the accuracy of the alignment (Supplementary figure 4D and Supplementary table 1). As described in Supplementary note 5 of Paul-Gilloteaux et al.,[3] the TRE at any spatial point depends on the distance of this point to the set of landmarks used for registration. Because all landmarks were placed in the vicinity of the mineral, the TRE ranged from 63 to 71 nm near the mineral and increased with distance, reaching 164, 287, and 623 nm in the XY, XZ, and YZ views, respectively. Since the TRE reflects the accuracy of the alignment, we consider an alignment to be acceptable when the maximum TRE is comparable to the resolution of the lowest-resolution imaging modality used (in this case, light microscopy), which is satisfied here.

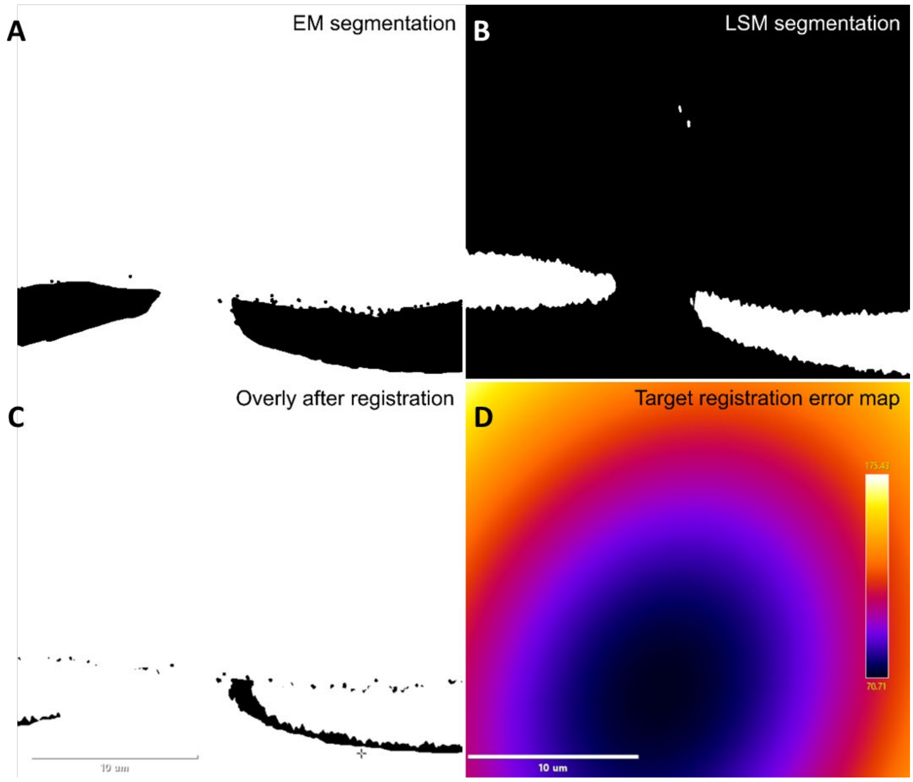


**Supplementary figure 4**. Example from the XY view showing (A) manual segmentation of an EM image, (B) segmentation of the corresponding LSM image, (C) overlay of the EM and LSM segmentations following registration using ec-CLEM, and (D) the predicted target registration error heat map generated by ec-CLEM, indicating errors between 63 and 623 nm for this view.

**Supplementary Table 1. Minimum and maximum target registration error values for all analyzed mini-stacks.**

| Mini stacks (view and slice numbers) | Minimum TRE (nm) | Maximum TRE (nm) |
|---|---|---|
| XY 1-5 | 63.25 | 171.09 |
| XY 20-25 | 63.00 | 156.60 |
| XY 50-55 | 63.25 | 171.45 |
| XY 100-105 | 63.25 | 147.79 |
| XY 150-155 | 63.25 | 162.55 |
| XY 200-205 | 70.71 | 175.43 |
| Average XY | 64.45 | 164.15 |
| XZ_450-455 | 63.25 | 236.53 |
| XZ_500-505 | 66.67 | 151.65 |
| XZ_550-555 | 70.71 | 472.76 |
| Average XZ | 66.87 | 286.98 |
| YZ_350-355 | 70.71 | 709.45 |
| YZ_450-455 | 70.71 | 730.58 |
| YZ_550-555 | 63.25 | 416.82 |
| YZ_600-605 | 75.59 | 637.73 |
| Average YZ | 70.65 | 623.65 |

**Supplementary figure 5: Thickness mesh from volumeSEM imaging**

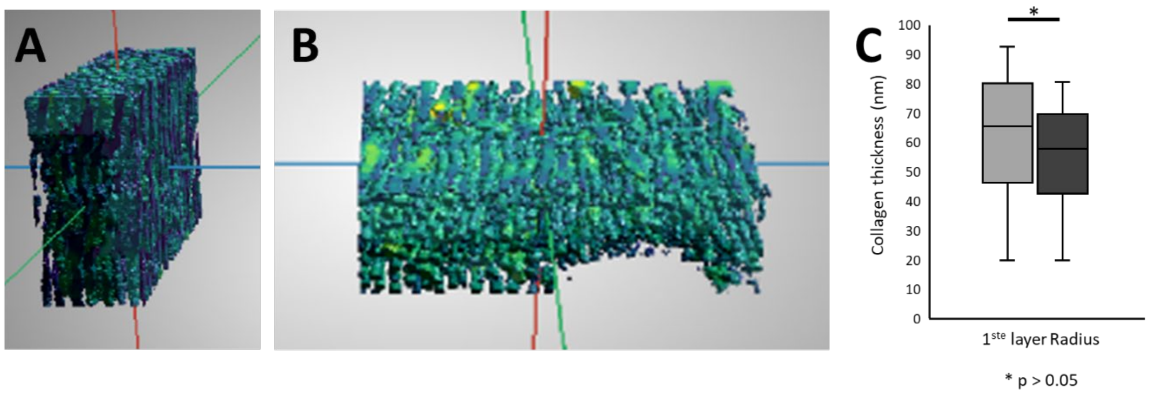


**Supplementary figure 5**: A, B Thickness mesh of selected regions from the segmented cryo-FIB/SEM images within the radius. C: results of the thickness measurements showing a significant difference between the collagen of the first continuous layer (grey) and the collagen within the radius (black).

**Supplementary figure 6: Live-to–cryo fluorescence microscopy: catching the moment AND the place**

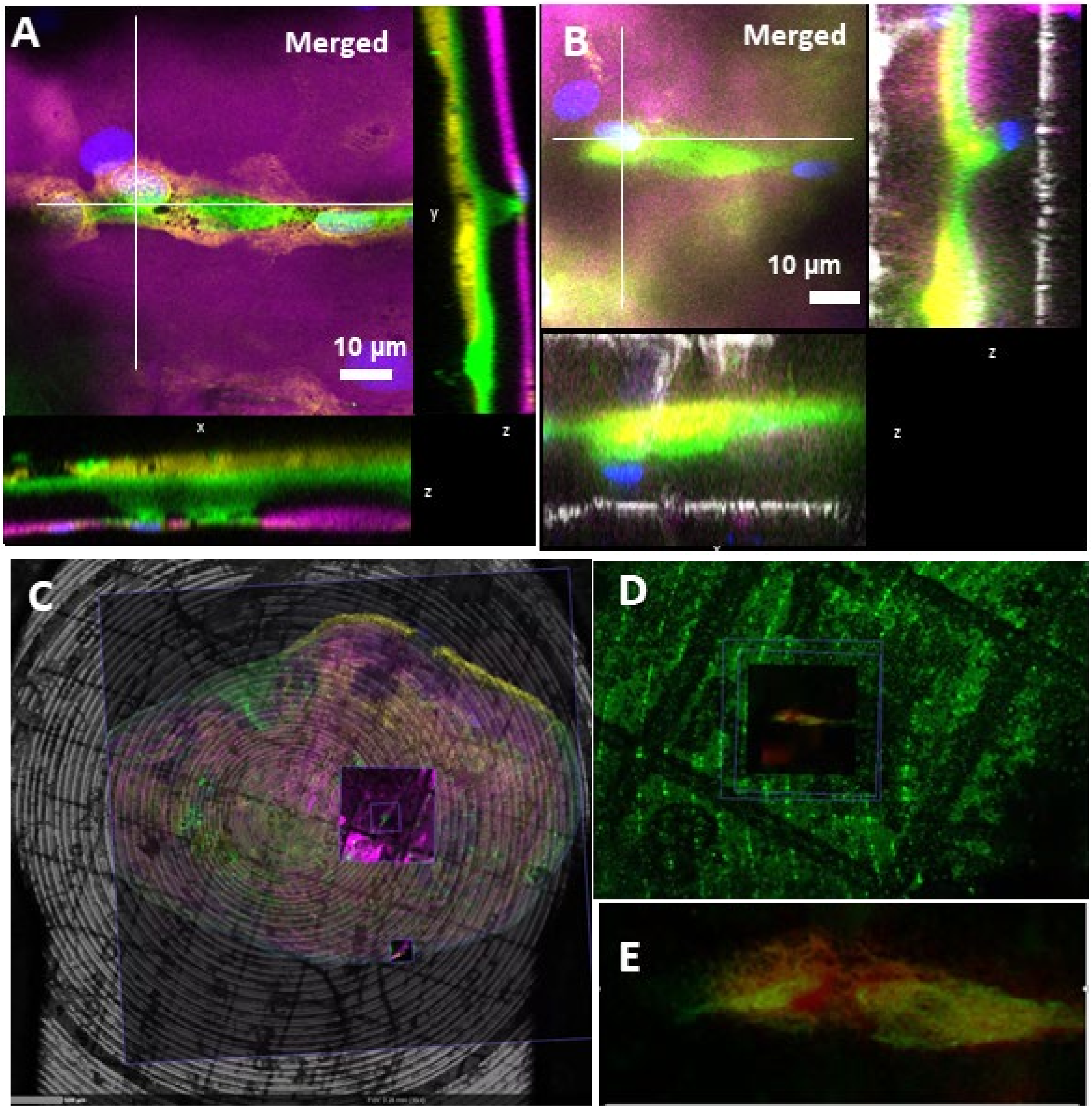


***Supplementary figure 6*** *Live-to-cryo fluorescence microscopy: A) Orthogonal view of a 63x magnification fluorescent 3D image of the 6 day old scale. Stained for elasmoblasts (yellow), mineral, purple, collagen (green) and nuclei (blue). B) orthogonal view of a 100x magnification cryo-fluorescent image of the same region as panel d. Colors are identical, white is reflection image. C) Overlay image of live- and cryo-fluorescent images of a 6 day old scale, showing the correlative approach to locate the regions of interest. D) overlay of a 10 and 100x cryo-fluorescent image with a 63x live-fluorescent image to displaying the high resolution correlation of the two imaging modalities. E) Overlay of a 63x live (green) with the 100x cryo (red) elasmoblast signal of the same cell shown in panel d, indicating perfect targeting of the ROI.*

**Supplementary figure 7: Preparation and correlation of TEM lamella from a cryo-preserved sample**

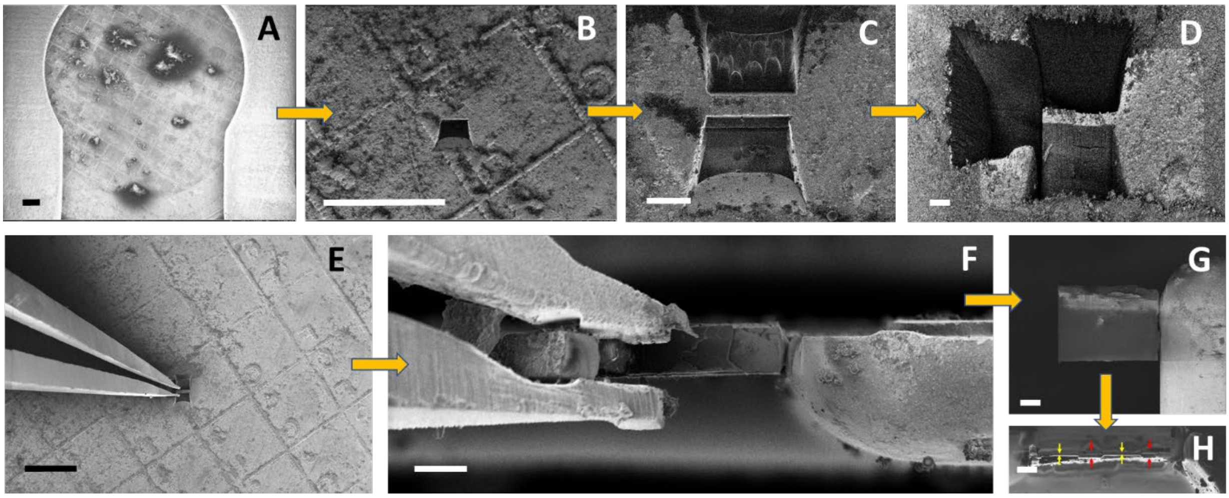


***Supplementary figure 7*** *Preparation and correlation of TEM lamella from a cryo-preserved sample. A) cryo-FIB overview of the surface of a cryo-preserved sample, showing the finder top pattern to allow for accurate localization. B) formation of the first trench approaching the TEM ROI at the required location, identified by the finder top-pattern after image correlation. C) completion of the second trench around the TEM ROI. D) completion of the third trench around the TEM ROI allowing for enough space to approach the sample with the gripper. E) approach of the gripper (Kleindiek, Germany), to the prepared lamella in the cryo-preseverd sample used to extract the ROI. F) attachment of the thick lamella to a half-moon grid (Aurion, the Netherlands), by use of the gripper in cryo-conditions (top view). G) Attached thick lamella to the half-moon TEM grid. H) Thinned TEM lamella with electron transparent windows (yellow arrows) suspended between thicker regions for stability (red arrows). Scale bars: a,b,e 300 µm; c,d,f,g,h 20 µm.*

**Supplementary figure 8 Fibril thickness determined in TEM lamella**

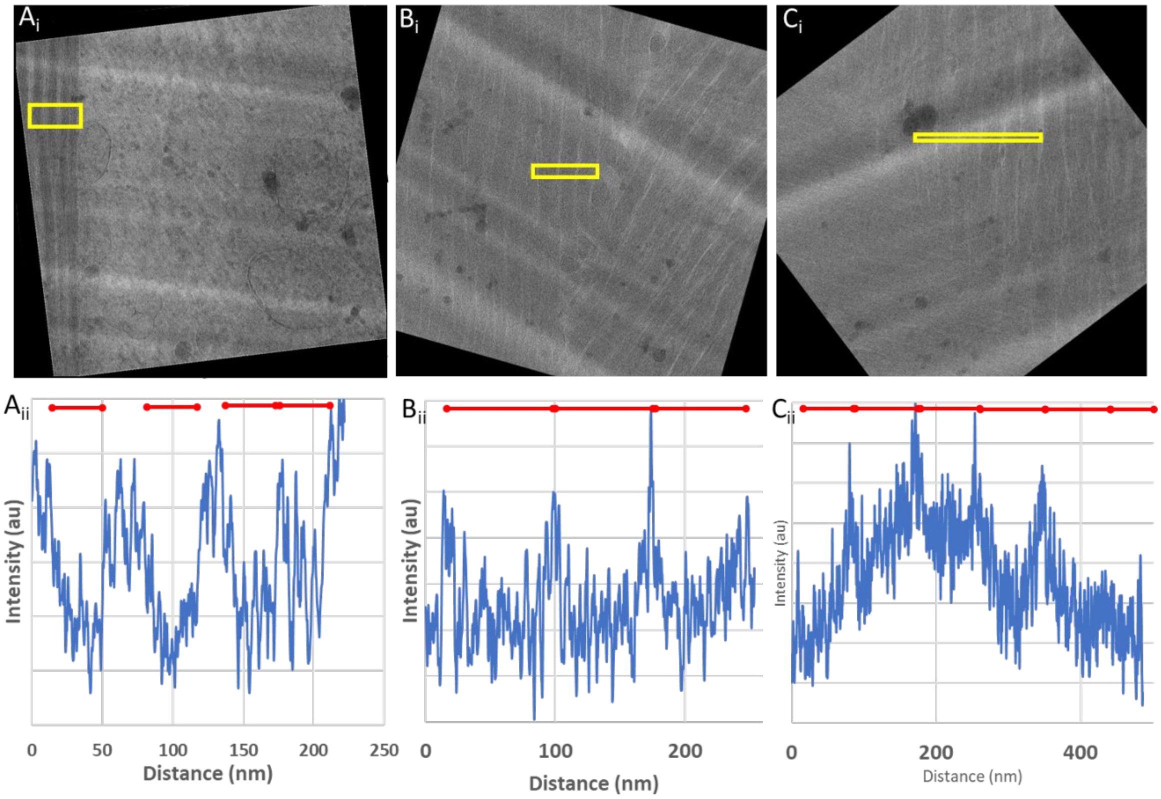


***Supplementary figure 8:*** *Fibril thickness determined in TEM lamella. A) Thickness measurement of collagen fibers close to the elamoblast, (region shown in Fig. 3 Ciii). B) Thickness measurement of collagen fibers at the stained collagen, (region shown in Fig. 3 Cii C) Thickness measurement of collagen fibers at the stained collagen, (region shown in Fig. 3 Cii). i: TEM image rotated for vertical alignment of the fibers, yellow box indicates the evaluated region. ii: intensity plot measured in the indicated region using ImageJ. Low intensity regions are used to determine fibril thickness, as indicated by the red lines above.*

**Supplementary Figure 9: deconvolution of the Raman mineral spectra**

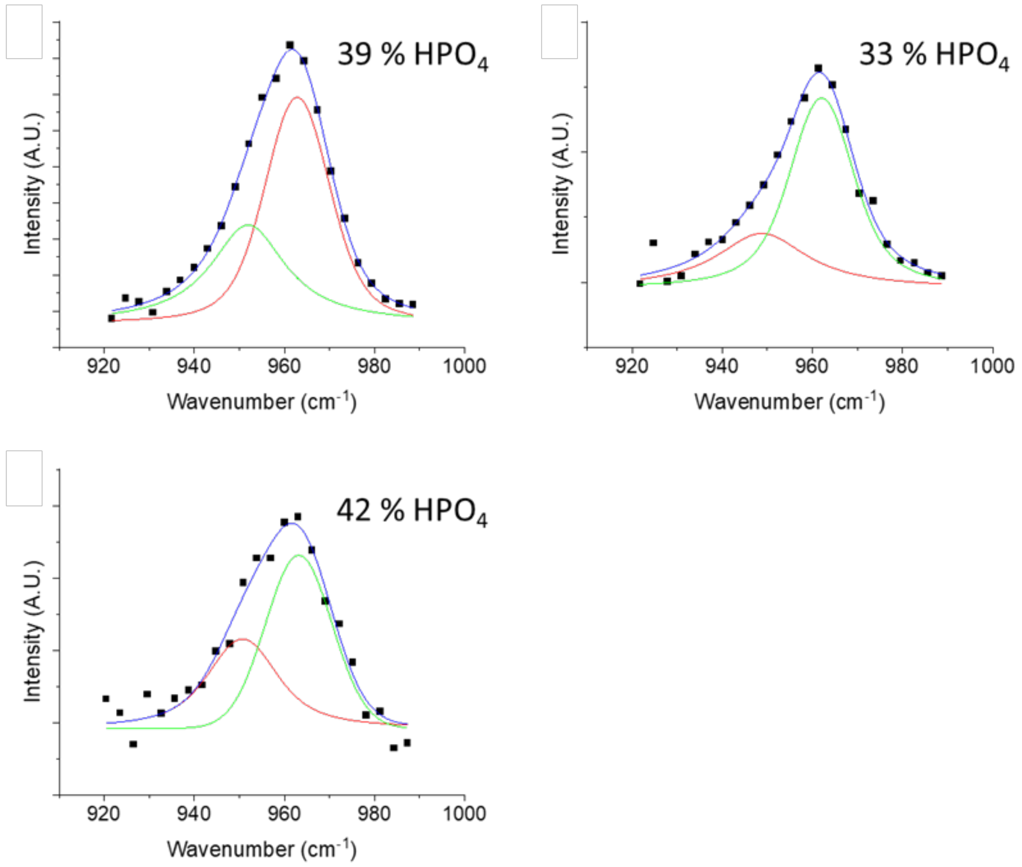


***Supplementary figure 9:*** *Deconvolution of the mineral in cryoRaman from zebrafish scales. 3 cryogenic zebrafish measurements (on lamella) deconvoluted for peaks for* $HPO_4$ *(950* $cm^{-1}$*) and* $PO_4$ *(962* $cm^{-1}$*). The percentage indicates amount of* $HPO_4$ *relative to total amount of total* $PO_4$ *(*$HPO_4$ *+* $PO_4$*).*

**Supplementary Fig. 10 Determination of the mineral composition**

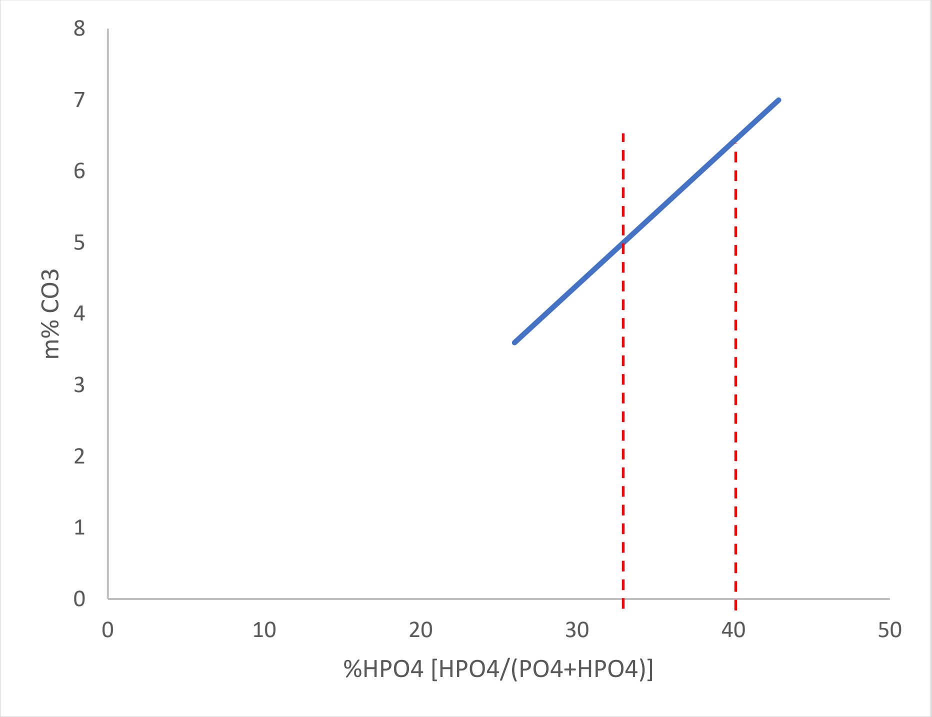


***Supplementary Fig. 10*** *Determination of the molecular formula ($Ca_{(10-0.5y)}$ $(PO_4)_{6-y-0.6x(10-0.5y)}$ $(CO_3)_y$ $(OH)_{2-0.2x(10-0.5y)}$ $(HPO_4)_{x(10-0.5y)}$) when carbonate and acidic phosphate are incorporated into the mineral. Blue line shows Relationship between carbonate substitution and phosphate when the molecular formula corresponds to the Ca/P ratio of 1.61 as determined by EDX. Red lines indicate the range of acidic phosphate incorporation determined with Raman microspectroscopy as shown in Supplementary Fig. 9.*

**All supplementary videos are available in the open access online repository Zenodo[4].**

**Supplementary video 1**
Video of the correlation from live-LSM to cryoFIB/SEM

**Supplementary video 2**
3D visualization of the radius as shown with cryoFIB/SEM combined with the correlation with the live-LSM imaging.

**Supplementary video 3**
3D-recreation of the cryoFIB/SEM stack groing through the different layers showing them to consist of aligned collagen fibers

**Supplementary video 4**
3D-recreation of a high resolution stack going through 2 layers of the collagen within the radius

**Supplementary video 5**
3D-recreation of a segmentation of the high resolution stack going through 2 layers of the collagen within the radius, as shown in SM 4.

**Supplementary video 6**
Movie showing the correlation from live-LSM to cryoTEM/tomography.

**Supplementary video 7**
Tomography of the TEM lamella of the collagen layers including the reconstruction and the segmentation of the collagen fibrils

**Supplementary video 8**
Tomography of the TEM lamella of the mineral layers including the reconstruction and the segmentation of the mineral platelets
